# Worldwide analysis of reef surveys sorts coral taxa by associations with recent and past heat stress

**DOI:** 10.1101/2021.10.29.466412

**Authors:** Oliver Selmoni, Gaël Lecellier, Véronique Berteaux-Lecellier, Stéphane Joost

## Abstract

Coral reefs around the world are under threat from anomalous heat waves that are causing the widespread decline of hard corals. Different coral taxa are known to have different sensitivities to heat, although variation in susceptibilities have also been observed within the same species living in different environments. Characterizing such taxa-specific variations is key to enforcing efficient reef conservation strategies.

Here, we combine worldwide-reef-survey data with remote sensed environmental variables to evaluate how local differences in taxa-specific coral cover are associated with past trends of thermal anomalies, as well as of non-heat related conditions. While the association with non-heat related environmental variation was seldom significant, we found that heat stress trends matched local differences in coral cover. Coral taxa were sorted based on the different patterns of associations with heat stress.

For branching, tabular and corymbose Acroporidae, reefs exposed to recent heat stress (measured the year before the survey) had lower coral cover than locally expected; and among these reefs, those previously exposed to frequent past heat stress (measured since 1985) displayed relatively higher coral cover, compared to those less frequently exposed. For massive and encrusting Poritidae, and for meandroid Favidae and Mussidae, we observed a negative association of coral cover with recent heat stress; however, unlike with Acroporidae, these associations were weaker and did not vary with past heat exposure. For Pocilloporidae, we found a positive association between coral cover and recent heat stress for reefs frequently exposed to past heat, while we found a negative association at reefs less frequently exposed to past heat. A similar pattern was observed for the branching Poritidae, although the associations were weaker and not statistically significant.

Overall, these results show taxa-specific heat association patterns that might correspond to taxa-specific responses to past heat exposure, such as shifts in the assembly of coral communities, evolutionary adaptation or physiological acclimation.

## 1 Introduction

For over 30 years now, coral reefs around the world have been suffering from a widespread decline of hard corals (Ateweberhan et al., 2011; De’ath et al., 2012; Cramer et al., 2020). This decline threatens the persistence of entire coral reef ecosystems, as hard corals constitute the physical architecture of such habitats. A leading cause of this decline is coral bleaching, a stress response driven by heat stress that causes corals to separate from the symbiotic algae living in their tissues (van Oppen and Lough, 2009). If this persists over several weeks, a bleaching state will eventually lead to coral death (Diaz-Pulido and McCook, 2002). Extensive reef deterioration following anomalous heat waves have been observed globally, with an estimated 3.2% loss of absolute coral cover (i.e. the percentage of the reef surface covered by live stony coral) worldwide from 2005 to 2015 (Souter et al., 2021). By 2050, it is expected that coral bleaching conditions will become persistent worldwide (van Hooidonk et al., 2013).

The sensitivity to thermal stress varies between coral species (Marshall and Baird, 2000; Loya et al., 2001; McClanahan et al., 2001). Growth form has typically been considered as one of the main proxies to assess heat stress susceptibility, where corals with branching morphologies were observed to be more sensitive to heat stress than corals with encrusting or massive morphologies (Loya et al., 2001). Coral growth form is correlated with several physiological and metabolic traits (e.g. tissue thickness, fecundity, growth rate), and combination of such traits were employed to define groups of corals with different life history strategies: such as “stress tolerant”, “generalist”, “weedy” and “competitive” (Darling et al., 2012). As reefs are hit by heat waves, their coral assemblies are shifting toward stress tolerant species (Hughes et al., 2018). Of note, substantial differences in heat tolerance can also be observed between coral of the same species (Bay and Palumbi, 2014; Schoepf et al., 2015; Louis et al., 2016; Klepac and Barshis, 2020; McClanahan et al., 2020b). Such differences can be due to the presence of genetic traits conferring heat tolerance to some colonies of a population (evolutionary adaptation), or to some colonies transiently adjusting their metabolism to cope with a stressful condition (physiological acclimation; Palumbi et al., 2014). Characterizing how responses to past heat exposure (*i.e*. community shifts, adaptation, acclimation) vary between taxa is of paramount importance in order to organize effective conservation efforts (Baums et al., 2019; Matz et al., 2020).

Over the last decade, the combination of field survey records with remote sensed data for sea surface temperature has become one of the major tools used to measure coral decline driven by heat stress. Multiple studies have shown associations between heat stress events and coral cover decline at both local (Head et al., 2019; Babcock et al., 2020; Selmoni et al., 2020a) and global scales (Selig et al., 2012). In other studies, the bleaching intensity metric was found to be positively associated with heat stress on the Australian Great Barrier Reef (Hughes et al., 2018, 2019), as well as across larger spatial scales (for instance, the Indo-Pacific region; Sully et al., 2019; McClanahan et al., 2020a). Notably, some of these studies investigated how past and recent thermal stress interact to drive coral cover decline/bleaching intensity, suggesting the existence of changes in thermal tolerance driven by past heat exposure (Thompson and van Woesik, 2009; Guest et al., 2012; Hughes et al., 2018, 2019; Head et al., 2019; Sully et al., 2019; McClanahan et al., 2020a; Selmoni et al., 2020a). Yet, few of these works focused on the differences in heat stress responses between coral taxa (Guest et al., 2012; Hughes et al., 2018; Head et al., 2019), and even fewer investigated these responses across wide spatial scales (e.g. the entire Indo-Pacific, McClanahan et al., 2020a). Of note, previous studies detected a strong spatial variability in coral heat responses, and suggested that the effects of additional variables (e.g. relating to temperature variability, as well as environmental constraints such as water velocity, sedimentation, eutrophication) might confound or interact with heat stress (Maina et al., 2011; Safaie et al., 2018; McClanahan et al., 2019, 2020a, 2020b; Sully et al., 2022).

In the present study, we combine pre-existing global coral cover data obtained from the Catlin Seaview Survey project (González-Rivero et al., 2014, 2016) with time series of sea surface temperature anomalies computed from remote sensed data by the Coral Reef Watch (Skirving et al., 2020). The goal is to investigate how spatial patterns of historical heat stress overlap with local differences in coral cover of different coral taxa. We first evaluate how long-lasting trends of heat stress and alternative environmental constraints (related for instance to water turbidity, salinity and current velocity) are associated with local differences in coral cover. These differences concern (1) overall coral cover, and (2) coral cover specific to seven taxa whose distribution spans across different oceans. Next, we assess how period-specific trends of heat stress associate with local differences in coral cover, and in particular how such associations with recent heat stress (measured across the year before the survey) interacts with (i.e. are accentuated or mitigated by) past heat stress exposure (measured during all the previous years since 1985). The results highlight four groups of coral taxa showing distinct patterns of heat associations.

## 2 Materials and methods

### 2.1 Coral cover data

The coral cover data were obtained from the Catlin Seaview Survey (CSS) project (González-Rivero et al., 2014, 2016). When we accessed the data in January 2021, there were a total of 860 surveys performed along 579 transects (380 transects were visited once, 140 twice, 36 three times, and 23 four times) across twelve study areas around the world between 2012 and 2018 (Figure 1A). The CSS project applied the same standardized framework to each survey when recording and analyzing field data. First, field surveys were performed using an underwater 360° camera that took a picture every three seconds along a 1.6-2 km transect, maintaining a constant depth of 10 meters. Next, field pictures were processed using automated image recognition based on machine learning algorithms. Such algorithms were trained using labels that were manually annotated by coral taxonomists (Beijbom et al., 2012).

**Figure 1.**
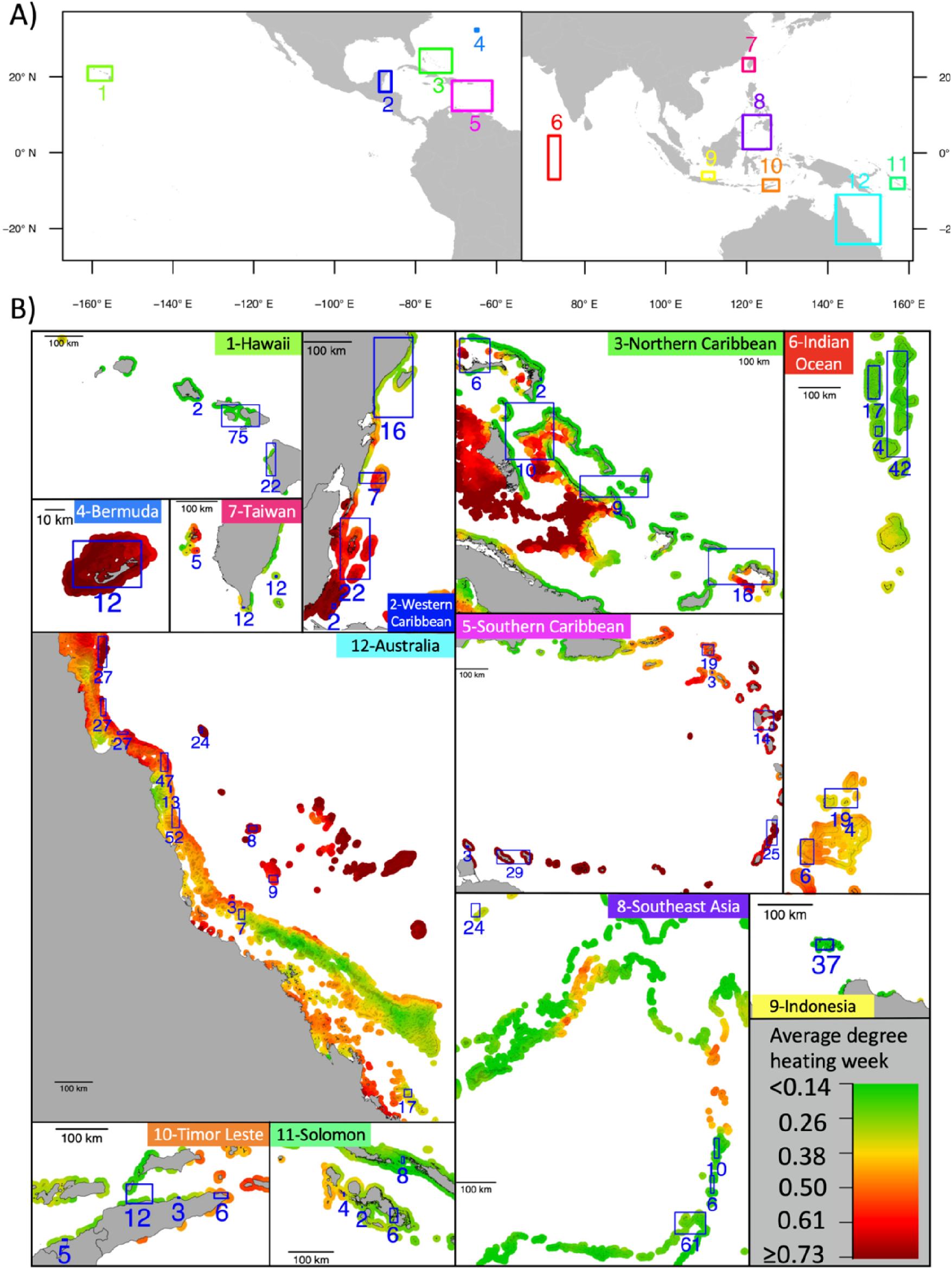
The study regions. A) Distribution of the twelve study regions. B) For each study region, the map shows the average monthly maximal Degree Heating Week (DHW) values for the 1985-2020 period. The blue squares indicate areas where coral cover surveys from the Catlin Seaview Surveys project were performed, with the numbers corresponding to the number of surveys per area. The reef surfaces were derived from the “Global Distribution of Coral Reefs” dataset of the UNEP (UNEP-WCMC et al., 2021).

For each reef survey, the CSS data provided (1) the overall measure of hard coral cover and (2) coral cover for 35 taxa of corals labelled using morphological characteristics (e.g. branching Acroporidae, massive Poritidae, meandroid Favidae and Mussidae). We used these data to obtain taxa-specific measurement of coral cover. As the number of labels employed varied between regions, and different regions featured different numbers of surveys, we focused only labels that appeared in at least half of the twelve study regions. After this filtering step, the dataset included taxa-specific coral cover data with the following seven labels: branching Acroporidae (CSS label ACR.BRA), tabular, corymbose, digitate Acroporidae (ACR.TCD), Meandroid Favidae and Mussidae (FAV.MUS), Pocilloporidae (POCI), massive Poritidae (POR.MASS), encrusting Poritidae (POR.ENC), branching Poritidae (POR.BRA).

### 2.2 Environmental data

Environmental data characterizing heat stress data were retrieved from the Coral Reef Watch database, as part of the 5 km-resolution sea surface temperature products (Skirving et al., 2020). The variable we used to describe heat stress was the Degree Heating Week (DHW, Figure 1B), as it has been shown to be directly correlated with coral bleaching occurrence and severity (Liu et al., 2014). For a given day, DHW is calculated as the sum of the temperature hotspots (that is, daily temperatures that exceeded the maximal monthly average by 1°C) from the preceding 12-weeks period. We used the CSS metadata to retrieve the coordinates of every survey location (defined by the transect mid-points) and at each coordinate we extracted the monthly maximal DHW from 1985 to 2020 using the RASTER R package (v. 3.0; Hijmans, 2021).

Using the same methods, we extracted the monthly averages for different datasets describing seascape conditions that potentially could be associated with changes in coral cover at survey sites, including:

1. chlorophyll concentration (CHL), accessed from the Copernicus Marine Services database (product id: OCEANCOLOUR_GLO_CHL_L4_REP_OBSERVATIONS_009_082, spatial resolution: 4 km, temporal window: 1997-2020, EU Copernicus Marine Service, 2017);
2. sea current velocity (SCV), accessed from the Copernicus Marine Services database (product id: GLOBAL_REANALYSIS_PHY_001_030, spatial resolution: ~8 km, temporal window: 1993-2020, EU Copernicus Marine Service, 2017);
3. suspended particulate matter (SPM), accessed from the Copernicus Marine Services database (product id: OCEANCOLOUR_GLO_OPTICS_L4_REP_OBSERVATIONS_009_081, spatial resolution: 4 km, temporal window: 1997-2020, EU Copernicus Marine Service, 2017);
4. sea surface salinity (SSS), accessed from the Copernicus Marine Services database (product id: GLOBAL_REANALYSIS_PHY_001_030, spatial resolution: ~8 km, temporal window: 1993-2020, EU Copernicus Marine Service, 2017);
5. sea surface temperature (SST), accessed from the Coral Reef Watch database (product id: ct5km_sst-mean_v3.1, spatial resolution 5 km, temporal window: 1985-2020; (Skirving et al., 2020).

For each of the six datasets (DHW, CHL, SCV, SPM, SSS and SST) used to describe seascape conditions at the CSS survey locations, we computed three statistics summarizing long-lasting environmental trends: mean (*VAR_MEAN_*), standard deviation (*VAR_STD_*) and maximal value (*VAR_MAX_*). The long-lasting trends covered the entire temporal window from the first measurement of the environmental variable until the date of the CSS survey.

For DHW, we computed twelve additional variables that decompose overall trends into period-specific trends of heat stress. These twelve variables were computed using three different statistics: mean (*DHW*_*MEAN*_…_), standard deviation (*DHW*_*STD*_…_) and maximal value (*DHW*_*MAX*_…_), each across four different time periods (Figure 2). The first period represents recent heat stress and covers the year that preceded the survey date (*DHW*_…_1*yr*_; Figure 2a). Similarly, we defined past heat stress as the periods covering the 10 years (*DHW*_…_10*yrs*_; Figure 2b) and the 20 years (*DHW*_…_20*yrs*_; Figure 2c) that preceded the survey. Additionally, we defined one long-term variable of past heat stress which excluded the first year before the sampling date and covered all the previous years (*DHW*_…_>1*yr*_; Figure 2d). This variable was developed as an estimator of past heat stress that does not overlap with recent heat stress, so that it could be used in bi-variate model construction (see the “Statistical analysis” section).

**Figure 2.**
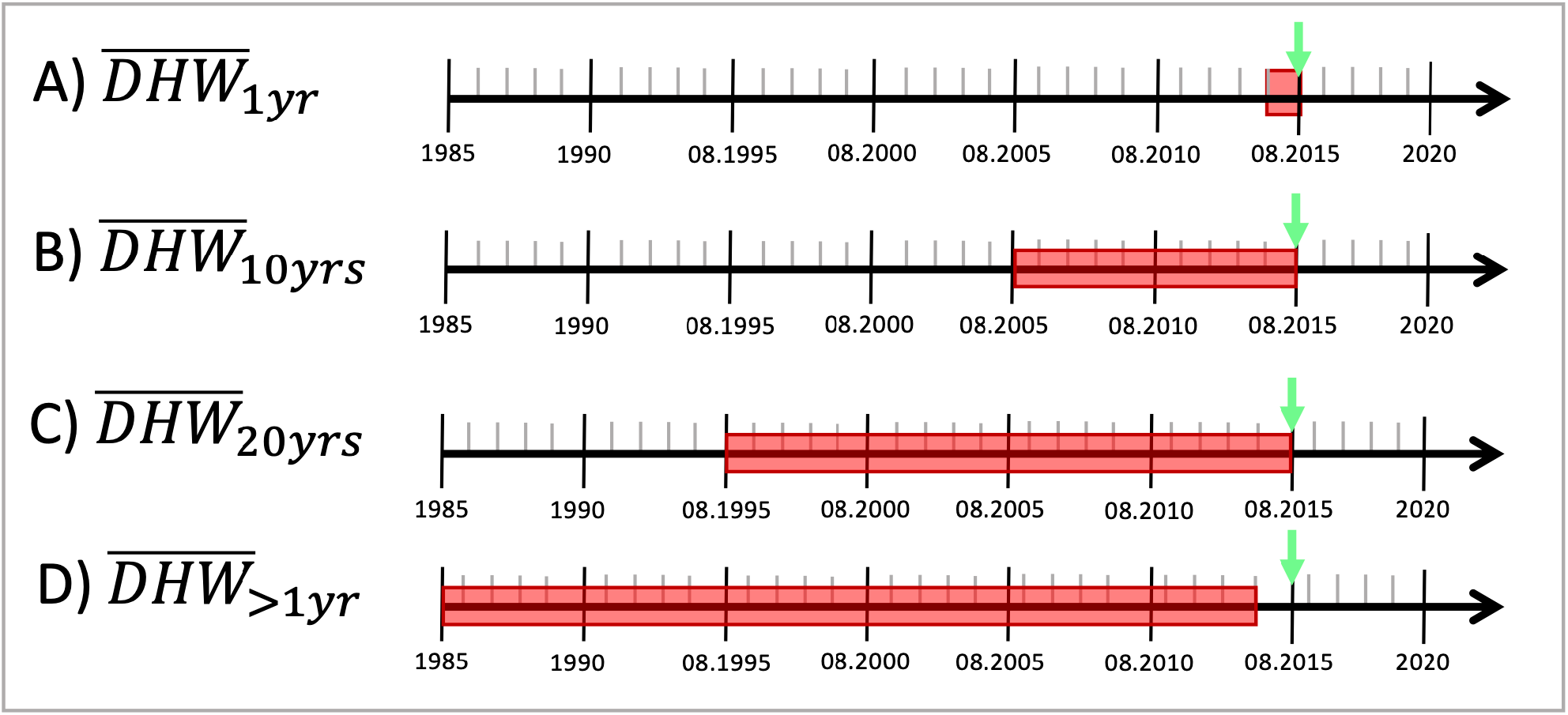
Degree heating week (DHW) periods. The figure summarizes the four temporal windows across which three period-specific DHW statistics (mean, standard deviation, maximal value) were calculated. The timelines are hypothetical examples, where the green arrows indicate a hypothetical date of the coral cover survey, and the red boxes show the periods of interest used to calculate period-specific DHW statistics.

Hereafter, the variable names are abbreviated using the dataset name, followed by the statistics’ name and the time period concerned (i.e., *DHW_statistics_period_*). For instance, *DHW*_*STD*_10*yrs*_ is the standard deviation of monthly maximum DHW measured across the 120 months (10 years) that preceded the survey date.

### 2.3 Statistical analysis

Previous studies have employed repeated measures of coral cover over time to determine the trends of coral growth or decline, and then used statistical analyses to match these trends with environmental data (Selig et al., 2012; Head et al., 2019; Babcock et al., 2020). This approach is not transposable to the CSS dataset, as most of the transects were surveyed only once. For this reason, we employed a different method to investigate how local differences in coral cover were associated with patterns of environmental variation measured at the survey sites.

In practice, we investigated the association between coral cover and environmental variation using generalized linear mixed-models (GLMMs), where the random factors controlled for spatial autocorrelation between survey sites (Dormann et al., 2007). We employed three levels of random factors representing spatial autocorrelation at different spatial scales and progressively nested into each other (i.e. the factors covering smaller spatial scales were nested inside those covering larger spatial scales). The first random factor was the study area (twelve levels, Figure 1A). We then designed two additional random factors to control for spatial autocorrelation at a regional and at a local scale (within each study area). This was done by applying the following clustering approach in the R environment. First, we computed the Euclidean distances between surveys sites, and applied a hierarchical clustering using the Ward distance method. The result was a tree of distances between survey sites that was used to compute the two additional random factors: (1) at regional-level, survey sites located up to 100 km from each other; and (2) at local-level, survey sites located up to 7 km from each other. The two threshold values (100 km and 7 km) used in the clustering procedure were chosen as they corresponded to the average value and the minimal value, respectively, of the mean Euclidean distances measured between survey sites within each study area.

GLMMs were built using the glmmtmb R package (v. 1.0; Brooks et al., 2017). We investigated eight response variables recorded at each survey site: overall coral cover and taxa-specific coral cover for seven taxa described in the “Coral cover data” section (ACR.BRA, ACR.TCD, FAV.MUS, POCI, POR.MASS, POR.ENC, POR.BRA). As coral cover values are percentages, we used the beta regression method to model their response (Ferrari and Cribari-Neto, 2004). For each coral cover variable, we constructed three types of GLMMs (Table 1):

**Univariate GLMMs of long-lasting environmental trends:** the goal of these models is to assess how local patterns of coral cover can be explained by long-lasting trends (mean, standard deviation and maximal value) of variables describing distinct conditions: degree heating week (DHW; M1 to M3 in Table 1A), chlorophyll concentration (CHL: M4 to M6), sea current velocity (SCV: M7 to M9), suspended particulate matter (SPM: M10 to M12), sea surface salinity (SSS: M13 to M15) and sea surface temperature (SST: M16 to M18).
**Univariate GLMMs of period-specific trends of heat stress:** the goal of these univariate models is to evaluate how local patterns of coral cover can be explained by recent and past heat stress. We investigated the association of the coral cover variables with each of the nine explanatory variables describing averages, standard deviations and maximal values of DHW calculated across periods of 1 year (*DHW*_…_1*yr*_; M1 to M3 in Table 1B), 10 years (*DHW*_…_10*yrs*_; M4 to M6) or 20 years (*DHW*_…_20*yrs*_; M7 to M9) before the survey.
**Bi-variate GLMMs of modulating effects of past heat stress**: the goal of these bi-variate models is to evaluate how past heat stress interacts (i.e. accentuates or mitigates) with the association between recent heat stress and coral cover. In practice, the bi-variate models describe the associations between coral cover variables and two explanatory terms. The first term was the same for all bi-variate models. This term represents recent heat, and corresponds to the maximal DHW value measured during the year before the survey (*DHW*_*MAX*_1*yr*_). We chose this variable as it was the one showing the highest goodness-of-fit for most of the coral cover variables in univariate models (see Results). The second term corresponded to the interaction between (1) *DHW*_*MAX*_1*yr*_ and (2) each of the three explanatory variables describing average, standard deviation and maximum values of DHW calculated over all the previous years (*DHW*_…_>1*yr*_; M13 to M15). Importantly, the interaction variables describing long-term heat stress trends (*DHW*_…_>1*yr*_) were weakly collinear with *DHW*_*MAX*_1*yr*_ (R<0.2, Supplementary Figure 1).

**Table 1.** List of the models computed. The table displays the formulae of the univariate and bi-variate models computed for each coral cover variable (CC, representing either overall or taxa-specific coral cover). Table A) shows the formulae of models for univariate models based on long-lasting environmental trends (mean, standard deviation and maximal value) of degree heating week (DHW; M1-3), chlorophyll concentration (CHL; M4-6), sea current velocity (SCV; M7-9), suspended particulate matter (SPM; M10-12), sea surface salinity (SSS; M13-15) and sea surface temperature (SST; M16-18). Table B) displays the formulae of models based on period-specific trends (mean, standard deviation and maximal value) of DHW measured during the year (M19-21) the ten years (M22-24) and the twenty years (M25-27) before the survey. Table C) shows the formulae of bi-variate models based on the interaction between recent heat stress and past trends of heat stress. The first explanatory variable, representing recent heat stress, is always the maximal value of DHW measured during the year before the survey (*DHW*_*MAX*_1*yr*_). The second explanatory variable is the interaction of *DHW*_*MAX*_1*yr*_ with the past trends (averages, standard deviations and maximal values) of DHW measured during all of the previous years (*DHW*_…_>1*yr*_; M28-30). *α* indicates the intercept, *β* the effects of the explanatory variables.

| A) Univariate models of long-lasting environmental trends |  |  |
| --- | --- | --- |
| *Degree heating week - DHW* | *Sea current velocity - SCV* | *Sea surface salinity - SSS* |
| M1) $CC \sim \alpha + \beta(DHW_{MEAN})$ | M7) $CC \sim \alpha + \beta(SCV_{MEAN})$ | M13) $CC \sim \alpha + \beta(SSS_{MEAN})$ |
| M2) $CC \sim \alpha + \beta(DHW_{STD})$ | M8) $CC \sim \alpha + \beta(SCV_{STD})$ | M14) $CC \sim \alpha + \beta(SSS_{STD})$ |
| M3) $CC \sim \alpha + \beta(DHW_{MAX})$ | M9) $CC \sim \alpha + \beta(SCV_{MAX})$ | M15) $CC \sim \alpha + \beta(SSS_{MAX})$ |
| *Chlorophyll concentration - CHL* | *Suspended particulate matter - SPM* | *Sea surface temperature - SST* |
| M4) $CC \sim \alpha + \beta(CHL_{MEAN})$ | M10) $CC \sim \alpha + \beta(SPM_{MEAN})$ | M16) $CC \sim \alpha + \beta(SST_{MEAN})$ |
| M5) $CC \sim \alpha + \beta(CHL_{STD})$ | M11) $CC \sim \alpha + \beta(SPM_{STD})$ | M17) $CC \sim \alpha + \beta(SST_{STD})$ |
| M6) $CC \sim \alpha + \beta(CHL_{MAX})$ | M12) $CC \sim \alpha + \beta(SPM_{MAX})$ | M18) $CC \sim \alpha + \beta(SST_{MAX})$ |
| B) Univariate models of period-specific heat stress trends |  |  |
| *1 year period* | *10 year period* | *20 year period* |
| M19) $CC \sim \alpha + \beta(DHW_{MEAN\_1yr})$ | M22) $CC \sim \alpha + \beta(DHW_{MEAN\_10yrs})$ | M25) $CC \sim \alpha + \beta(DHW_{MEAN\_20yrs})$ |
| M20) $CC \sim \alpha + \beta(DHW_{STD\_1yr})$ | M23) $CC \sim \alpha + \beta(DHW_{STD\_10yrs})$ | M26) $CC \sim \alpha + \beta(DHW_{STD\_20yrs})$ |
| M21) $CC \sim \alpha + \beta(DHW_{MAX\_1yr})$ | M24) $CC \sim \alpha + \beta(DHW_{MAX\_10yrs})$ | M27) $CC \sim \alpha + \beta(DHW_{MAX\_20yrs})$ |
| C) Bi-variate models of interactions between recent and past heat stress |  |  |
| M28) $CC \sim \alpha + \beta_1(DHW_{MAX\_1yr}) + \beta_2(DHW_{MAX\_1yr}:DHW_{MEAN\_>1yr})$ | | |
| M29) $CC \sim \alpha + \beta_1(DHW_{MAX\_1yr}) + \beta_2(DHW_{MAX\_1yr}:DHW_{STD\_>1yr})$ | | |
| M30) $CC \sim \alpha + \beta_1(DHW_{MAX\_1yr}) + \beta_2(DHW_{MAX\_1yr}:DHW_{MAX\_>1yr})$ | | |

For each GLMM, we reported the estimate (*β*, with its standard deviation) and the p-value from the Wald test for the fixed effects. We also computed an approximation of the estimate of the fixed effects (*β_resp_*) in the unit scale of the response variable (i.e. in coral cover percentage), using the R-package VISREG (v. 2.7; Breheny and Burchett, 2017) and custom R functions.

In addition, we evaluated the goodness-of-fit of each of the GLMMs by measuring the difference between the model’s Akaike Information Criterion (dAIC; Bozdogan, 1987) and the AIC of a null model. For univariate models, the corresponding null model was a GLMM using a constant value as explanatory variable. For bi-variate models, the null model was the univariate GLMMs using *DHW*_1*yr*_ as explanatory variable (M21). This approach allowed us to evaluate how accounting for the interaction with past heat stress improves the goodness-of-fit, compared to a model including recent heat stress only (*DHW*_1*yr*_). According to the rules of thumb for model selection (Burnham and Anderson, 2004), a model with *dAIC* < −2 (where *dAIC* = *AIC_null model_* – *AIC_model_*) has an improved goodness-of-fit, compared to the null model.

For each GLMM, we calculated the coefficients of variation (CV) related to the three random factors accounting for spatial autocorrelation (at study area-, regional- and local-level). CV was calculated by dividing the conditional standard deviation of every random factor by the intercept value of the model. In practice, these CVs could then be used as scaled standard deviations to compare the amount of variance controlled by every random factor across the different models (higher CV indicates larger amount of variance).

### 2.4 Grouping of taxa based on heat association

Based on the results of the association study between heat stress and taxa-specific coral cover, we categorized the seven coral taxa retained for the analysis into four groups showing distinct types of heat associations. The grouping was based on two criteria. First, whether differences in coral cover were significantly (*dAIC* < −2) associated with recent heat stress (*DHW*_*MAX*_1*yr*_) in the univariate model M21 (Table 1B). Second, whether past heat stress (*DHW_MEAN_>_1yr_*) interacted significantly (*dAIC* < −2) with recent heat stress (*DHW*_*MAX*_1*yr*_) in the bivariate model M28 (Table 1C). We chose M21 and M28 as these models display the highest goodness-of-fit in the association models with the majority of the coral cover variables (see Results). Taxa were assigned to the four heat association groups as follows:

Group 1 (GR1): significant association of local coral cover with recent heat, significant interaction with past heat.
Group 2 (GR2): significant association of local coral cover with recent heat, non-significant interaction with past heat.
Group 3 (GR3): non-significant association of local coral cover with recent heat, significant interaction with past heat.
Group 4 (GR4): non-significant association of local coral cover with recent heat, non-significant interaction with past heat.

For each survey, we computed the overall coral cover for every heat association group as the sum of the taxa-specific coral cover of taxa assigned to every group. Next, we computed the models of association with heat stress M21 and M28 for the overall coral cover of every heat association group using the same methods described in the “Statistical analysis” section. The goal of these models was to summarize average coral cover-heat associations for every heat association group. To visualize such average associations, we used the visreg R package (v. 2.7; Breheny and Burchett, 2017) and custom R functions.

## 3 Results

### 3.1 Associations with long-lasting environmental trends

Most of the models (16 out of 24 models) that used long-lasting DHW trends as explanatory variables (M1-M3) resulted in a stronger goodness-of-fit (*dAIC* < −2) when compared to a constant null model and displayed significant (p<0.01) negative associations (*β_resp_* < 0) with coral cover (Table 2A). These results were observed for models focusing on overall coral cover and for models focusing on taxa-specific coral cover of branching (ACR.BRA), corymbose, tabular and digitate Acroporidae (ACR.TCD), encrusting (POR.ENC) and massive (POR.MASS) Poritidae and meandroid Favidae and Mussidae (FAV.MUS). In contrast, models of Pocilloporidae (POCI) and branching Poritidae (POR.BRA) coral covers did not display significant associations with DHW trends. In general, models accounting for average DHW (M1) showed a weaker goodness-of-fit (higher values of *dAIC*), when compared with models based on maximal values of DHW (M3).

**Table 2.**
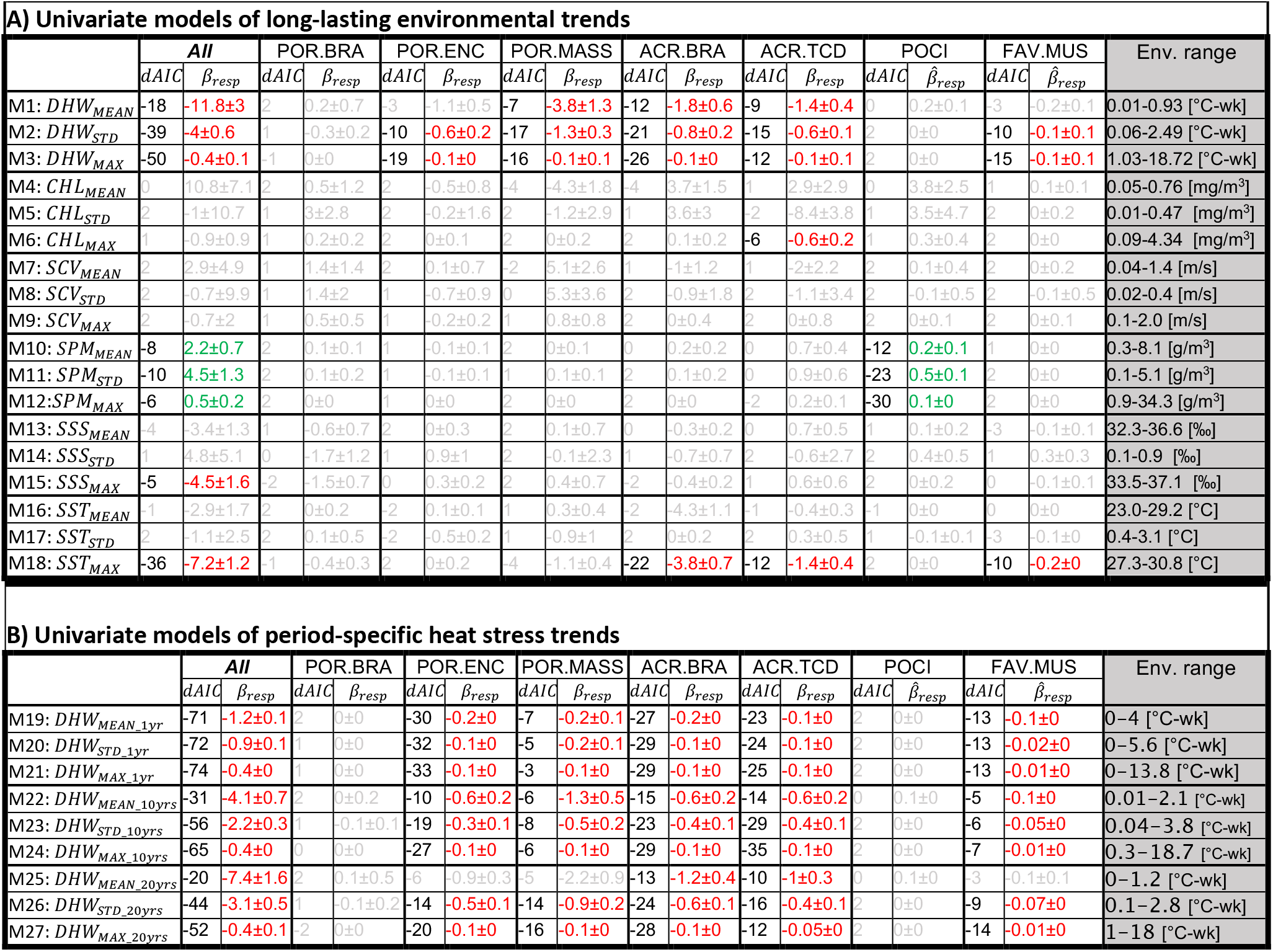
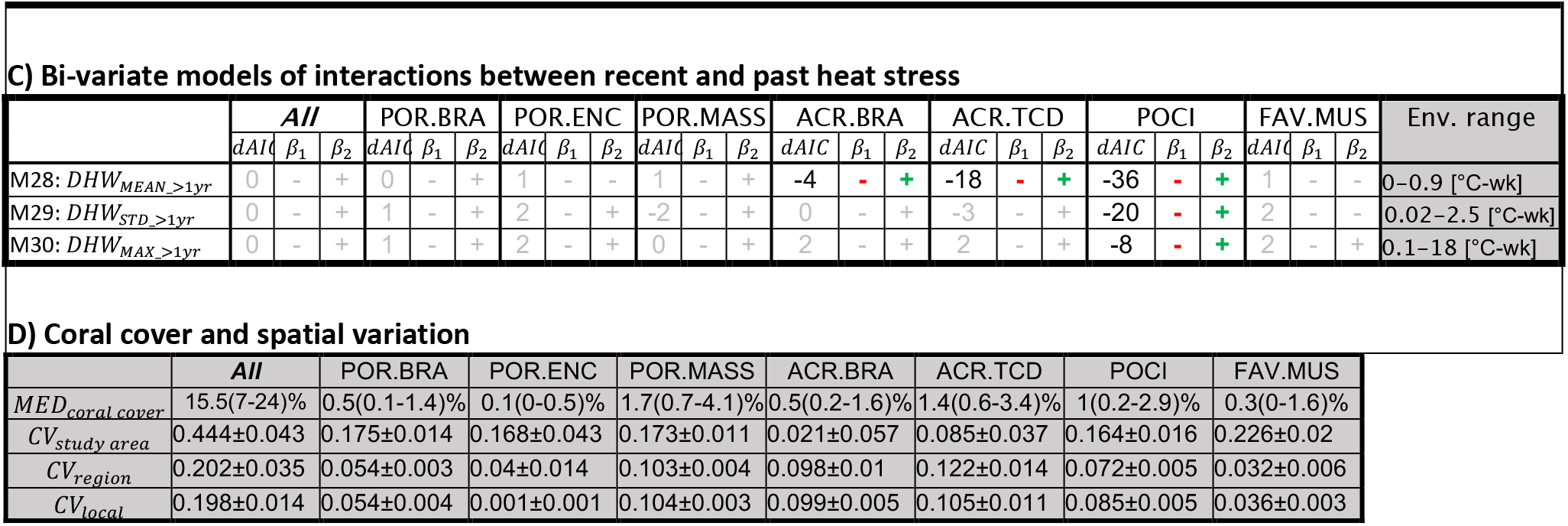
Association models of coral cover with environmental trends. The table shows the coefficients of models associated with different coral cover variables (columns) with variables describing environmental trends (rows). Coral cover variables refer to overall coral cover and taxa-specific coral cover (ACR.TCD= tabular, corymbose and digitate Acroporidae; ACR.BRA = branching Acroporidae; POR.ENC = encrusting Poritidae; POR.MASS = massive poritidae; FAV.MUS = meandroid Favidae and Mussidae; POCI = Pocilloporidae; POR.BRA = branching Poritidae), measured in field surveys. Environmental trends refer to the three statistics – mean, standard deviation and maximal value – applied to different variables: degree heating week (DHW), chlorophyll concentration (CHL), sea current velocity (SCV), suspended particulate matter (SPM), sea surface salinity (SSS) and sea surface temperature (SST). In table A), the models associate coral cover with long-lasting environmental trends, calculated across all the years prior to the survey date. In table B), models involve period-specific trends of DHW, measured over 1 year, 10 years or 20 years before the survey date. In table C), models involve two terms: (1) recent heat stress (maximal DHW measured during the year preceding the survey date) and (2) the interaction between recent heat stress and past heat stress (i.e. DHW statistics measured across all of the previous years). In table D), the median value (showing the interquartile range in parenthesis) of each coral cover variable (*MED_coral cover_*) is shown, together with the coefficients of variation associated with the three random factors controlling for spatial autocorrelation of the survey locations (at a study area-level, *CV_study area_*; at a regional-level, *CV_region_;* at a local-level, *CV_local_*). *dAIC* indicates the difference in the Akaike Information Criteria between each model and the null model (i.e. goodness-of-fit compared to the null model). In A) and B), *β_resp_* shows the estimated effect in the unit scale of response variable (i.e. the percentage of absolute coral cover; negative effects are in red and positive effects are in green). In C), *β*_1_ indicates the sign of the effect of recent heat stress on coral cover, and *β*_2_ indicates the sign of the effect of the interaction between recent and past heat stress. Cells in grey indicate models failing to show (1) a stronger goodness-of-fit when compared to the null model and (2) *β* significantly (p<0.01) different from 0. The last column headed ‘Env. range’ shows the range of the environmental variable implicated in the association model, with the units indicated in parentheses. The complete list of the coefficients and statistics describing the association models are displayed in the Supplementary Table 1.

For models employing explanatory variables other than DHW, we generally observed weaker goodness-of-fit and non-significant associations with coral cover variables (100 out of 112 models; Table 2). Among the few exceptions were the following significant associations: maximal CHL with coral cover of taxon ACR.TCD (M6 in Table 2A); SPM with overall coral cover and coral cover of taxon POCI (M10-12); maximal SSS with overall coral cover (M15); and maximal SST with overall coral cover and coral cover of taxa ACR.BRA, ACR.TCD and FAV.MUS (M18).

### 3.2 Associations with of period-specific trends of heat stress

Focusing on period-specific trends of heat stress, we observed that most models (52 out of 72 models) resulted in a stronger goodness-of-fit compared to a constant null model and displayed a significant negative association between DHW trends and coral cover (Table 2B). The goodness-of-fit generally appeared stronger in models based on DHW trends computed over shorter time periods (e.g. 1 year; M19-21 in Table 2B) and using the maximal value as the trend-statistic (M21, M24, M27). This was observed for overall coral cover and taxon-specific coral cover of taxa POR.ENC, ACR.BRA, ACR.TCD and FAV.MUS. Similar to patterns observed for long-lasting trends of DHW, no significant association with period-specific DHW trends were found for coral cover of taxa POR.BRA and POCI.

### 3.3 Interactions between recent and past heat stress

Regarding the bi-variate models, we observed that models accounting for the interaction of recent heat stress with average DHW trends over the long-term (M28) showed a stronger goodness-of-fit (when compared to the null models) for coral cover of taxa ACR.BRA, ACR.TCD and POCI. For these models, the association between recent heat stress and coral cover was of negative sign, and this negative association was contrasted by the interaction of positive sign with long-term heat stress. The same results were observed for coral cover of taxon POCI in models accounting for standard deviation and maximal DHW trends over the long-term (M29 and M30, respectively).

### 3.4 Sorting taxa by heat associations

We sorted taxa into four groups showing distinct types of heat associations (Figure 3).

**Figure 3.**
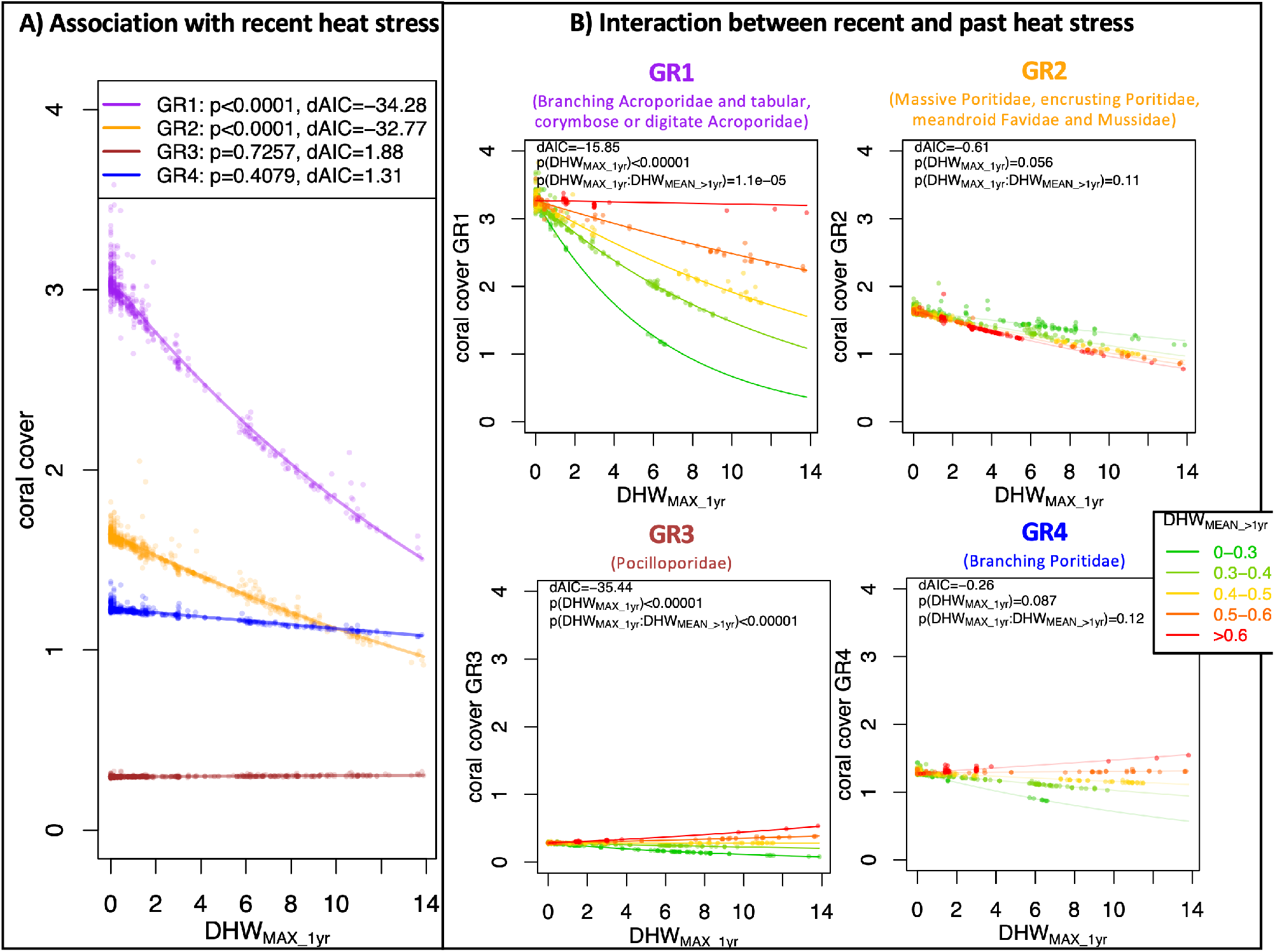
Groups of coral cover-heat stress associations. The plots display the associations between local coral cover and heat stress trends at survey sites for the four groupings of taxa with similar heat responses (heat association groups): in Group 1 (GR1, purple), taxa coral cover is associated with recent heat stress, where this association interacts with past heat stress; in Group 2 (GR2, orange), coral cover is associated with recent heat stress, without significant interaction with past heat stress; in Group 3 (GR3, brown), recent heat stress alone is not significantly associated with taxa coral cover, but the interaction between recent and past heat stress shows a significant association with coral cover; and in Group 4 (GR4, blue), no significant association between coral cover and heat stress was found. Plot A) displays the association of coral cover with recent heat stress, measured as the maximal degree heating week measured during the year before the survey (*DHW*_*MAX*_1*yr*_), for each of the four groups. Plots in B) show the association of coral cover with recent heat stress, for survey locations exposed to different levels of past heat stress, measured as the average DHW during all the previous years (*DHW*_*MEAN*_>1*yr*_). For each association model, the plots display the goodness-of-fit compared to a null model (*dAIC*) along with the p-values (p) of the fixed effects.

Group 1 (GR1) included ACR.BRA and ACR.TCD; these were the taxa that showed a negative association between coral cover and recent heat stress (*p* < 0.01; *dAIC* < −2), which was significantly modulated by the interaction with past heat stress. The association models for GR1 showed local differences in coral cover of −0.11 ± 0.02 % per °C-week of *DHW*_*MAX*_1*yr*_ (Figure 3A, purple regression line). In GR1, the local differences in coral cover associated with recent heat stress appeared to be mitigated at locations with higher past heat stress (Figure 3B, top-left graph). For example, a *DHW*_*MAX*_1*yr*_ of 1°C-week corresponded to a local difference in coral cover of −0.20 ± 0.03 % in locations where average past heat stress (*DHW*_*MEAN*_>1*yr*_) was below 0.3°C-week, whereas a difference in coral cover of −0.07 ± 0.01 % was observed in locations where average past heat was above 0.5°C-week.

Group 2 (GR2) included taxa that had a significant negative association between coral cover and recent heat stress, but without a significant modulation of the interaction with past heat stress. This group included the taxa POR.MASS, POR.ENC and FAV.MUS. The association model for GR2 showed local differences in coral cover of −0.05 ± 0.01 % per °C-week of *DHW*_*MAX*_1*yr*_ (Figure 3A, orange regression line). Accounting for the modulating interaction with past heat stress did not improve the goodness-of-fit (*dAIC* > −2) of the models when compared to the univariate counterpart accounting only for recent heat stress (Figure 3B, top-right graph).

Group 3 (GR3) included the POCI taxon alone. Here, significant differences in coral cover associated with recent heat stress were only detected when accounting for the modulating interaction with past heat stress. Indeed, the univariate model that only accounted for recent heat stress did not improve the goodness-of-fit (*dAIC* > −2) when compared to a constant null model (Figure 3A, brown regression line). However, the bi-variate model showed significant changes in the association between coral cover and recent heat stress, and such changes followed past heat stress levels (Figure 3B, bottom-left graph): when past heat stress was low, recent heat stress showed a negative association with coral cover, whereas when past heat stress was high, the association between coral cover and recent heat stress was of positive sign. For example, a *DHW*_*MAX*_1*yr*_ of 1°C-week corresponded to local differences in coral cover of −0.014 ± 0.002 % when average past heat stress (*DHW*_*MEAN*_>1*yr*_) was below 0.3°C-week, whereas it corresponded to local differences in coral cover of +0.007 ± 0.001 % when average past heat stress was above 0.5°C-week.

Group 4 included the POR.BRA taxon alone, where no significant differences in coral cover were detected in association with recent heat stress, nor with a modulating interaction of past heat stress (Figure 3A, blue regression line; Figure 3B, bottom-right graph).

## 4 Discussion

### 4.1 Local differences in coral cover mirror heat stress trends

We observed that survey sites exposed to elevated long-lasting and period-specific trends of Degree Heating Week (DHW) almost systematically displayed lower levels of coral cover than expected locally. These observations are consistent with previous studies where DHW-related variables were used as proxies for heat stress when investigating associations with coral decline or bleaching severity (Hughes et al., 2018; Head et al., 2019; McClanahan et al., 2019; Babcock et al., 2020). For the other environmental variables examined here (chlorophyll concentration, sea current velocity, suspended particulate matter, sea surface salinity and sea surface temperature), the associations with local differences in coral cover were generally weaker and seldom significant. As these environmental constraints were expected to drive local differences in coral cover based on past studies (Hédouin et al., 2015; Riegl et al., 2015; Jones et al., 2020; Sully and van Woesik, 2020), it is possible that the variables that we used are not appropriate proxies of such environmental constraints, particularly at the depth at which coral cover surveys were performed (see the “Limitations” section).

When we decomposed the effects of DHW into period-specific trends, we observed that association models with coral cover employing distinct DHW statistics (mean, standard deviation, and maximal value) as explanatory variables often displayed substantial differences in the goodness-of-fit. These results must be considered with care, due to the high collinearity (often > 0.8) between distinct DHW statistics measured during the same period (Supplementary Figure 1). Nevertheless, we note that univariate models employing DHW variables based on maximal values systematically provided a higher goodness-of-fit for the associations with coral cover, when compared to univariate models employing DHW averages or standard deviations. In the significant bi-variate models, we observed that reefs exposed to low levels of past heat stress showed a stronger negative association between coral cover and recent heat stress, compared to reefs exposed to higher levels of past heat stress. In contrast to univariate models, here this mitigating role of past heat was better explained by DHW averages than by DHW maximal values or DHW standard deviations. Average DHW might therefore represent the frequency of past heat stress that previous research found to be associated with decrease in bleaching rates (Thompson and van Woesik, 2009).

### 4.1 Taxon-specific heat associations

We classified coral taxa into four groups based on the patterns of coral cover association with recent and past heat stress.

Local differences in coral cover for taxa in Groups 1 (GR1) and 2 (GR2) matched the variation of recent heat stress, where reefs exposed to higher recent heat stress had lower coral cover than locally expected. The magnitude of this negative association was twice as strong for GR1 than for GR2. Assuming that these associations describe a causal relationship (i.e. recent heat stress drives local loss of coral cover), such differences between GR1 and GR2 might reflect differences in heat sensitivity. Indeed, GR1 includes heat sensitive taxa such as branching, corymbose and tabular *Acropora* and *Montipora*, while GR2 includes stress tolerant taxa such as massive and encrusting Poritidae, and meandroid Favidae and Mussidae (Loya et al., 2001; Darling et al., 2012; Guest et al., 2016; Hughes et al., 2018; Pisapia et al., 2019).

For coral taxa of heat association Groups 3 (GR3) and 4 (GR4), local differences in coral cover was not associated with variation in recent heat stress. However, previous studies listed taxa from these groups (GR3: Pocilloporidae, GR4: branching Poritidae) as “heat sensitive” (Loya et al., 2001; Guest et al., 2016) and “long-term losers” of coral bleaching (van Woesik et al., 2011). If the causal relationship assumed above were true, we expect that reefs exposed to higher recent heat stress would have lower coral cover for these taxa, compared to reefs exposed to lower recent heat. In fact, this negative association between coral cover of these taxa and recent heat stress was observed, but as we discuss below, it is only visible when considering the interaction with past heat stress.

When considered with the interaction with past heat stress, the association between coral cover and recent heat stress showed distinct patterns for the four heat association groups. For GR1, the negative association between coral cover and recent heat stress appeared to be mitigated by the interaction with past heat stress. This means that reefs exposed to high recent heat stress displayed lower coral cover than locally expected; however, among these reefs, those frequently exposed to past heat stress had relatively higher coral cover compared to reefs less frequently exposed. Of note, reefs exposed to very frequent past heat stress (*DHW*_*MEAN*_>1*yr*_ > 0.6°*C* – *week*) displayed similar coral cover regardless of recent heat stress exposure. GR3 had a similar, yet stronger, interaction as was observed for GR1. Indeed, in GR3 there was a negative association between coral cover and recent heat stress for reefs exposed to lower frequency of past heat stress, yet a positive association for reefs exposed to higher frequency of past heat stress. These results are consistent with previous local observations of taxa from these groups (*Acropora* and *Pocillopora*) that showed increased heat resistance after previous thermal exposure (Guest et al., 2012; McClanahan, 2017).

Similar increase in heat resistance was also previously observed for branching *Porites* (McClanahan, 2017). We found that GR4 (which includes branching Poritidae) showed similar associations and interactions to GR3, though these models were not found to be significant. The lack of clear statistical signal in these associations might be due heterogenous heat responses among species from GR4. For GR2 too, the interaction with past heat did not result as significant, but here the patterns were completely different from the other groups. Indeed, the interaction with past of heat was of negative sign, suggesting that past heat stress reinforced the negative association between recent heat stress and coral cover.

One possible explanation is based on the assumption that the associations between coral cover and heat stress are causal relationships, i.e. that recent heat stress exposure can drive local coral cover loss, and frequent exposure to past heat stress can provide a protection against such loss. Under this assumption, GR1, GR3 and some species in GR4 might have changed their thermal sensitivity in response to heat stress exposure over the past three decades. There are at least three (not mutually exclusive) phenomena that could underpin such rapid responses. The first phenomenon is a shift at the community level, where heat sensitive species have been progressively replaced by heat tolerant species from the same heat association group. This hypothesis assumes that there are substantial differences in heat tolerance between species from GR1 and GR3, which is possible even though corals within each of these groups usually have similar life-history traits and heat sensitivities (Loya et al., 2001; van Woesik et al., 2011; Darling et al., 2012). A second possible explanation is evolutionary adaptation, where colonies carrying genetic traits linked to thermal tolerance have been positively selected by recurrent exposure to heat stress. Indeed, recent environmental-genomics studies have suggested that the selection of such genetic traits could occur across relatively short time periods (~30 years) in both *Acropora* and *Pocillopora* species (Selmoni et al., 2020b, 2021). Nevertheless, this interpretation is hampered by the fact that every heat association group is composed of multiple species, and that each species likely has distinct adaptive potentials. The last phenomenon that could explain a hypothetical response to past heat exposure is physiological acclimation, where individuals exposed to past heat stress progressively adjust their metabolism, thus becoming more heat tolerant. However, acclimation is unlikely to play a key role here, as it usually occurs through constant heat stress exposure (e.g. large daily variation of SST; Palumbi et al., 2014), rather than via sporadic/seasonal heat exposure (i.e. decadal frequency of thermal anomalies).

### 4.2 Limitations

In this study, we investigated the associations between local coral cover and a set of environmental constraints measured with remote sensed data. This approach is inevitably exposed to bias because the spatial resolution of the environmental variables we employed is coarse (between 5 to 8 km), in comparison to the size of the transects providing the coral cover data (up to 2 km). The consequence of this mismatch in spatial resolutions is that habitat types (e.g., reef slope vs. reef flat) are overlooked by our analysis. Furthermore, the coral cover data that we employed was systematically collected at 10 meters of depth, such that our analyses might not be representative for coral cover at other depths.

Additionally, previous studies pointed out that geographic position is a key element for explaining variation in coral cover and bleaching responses (McClanahan et al., 2019; Sully et al., 2019). Our work is not an exception to this, as the random factors used to account for spatial autocorrelation among survey sites controlled for a substantial part of coral cover variation. This suggests that there are important factors proxied by geographical position that were not explicitly accounted for in our analyses. Such factors could be environmental conditions that we did not consider; or be environmental conditions that we did consider, but for which alternative types of descriptors should have be used. For instance, recent studies proposed measures of heat stress that are complementary to DHW, such as statistics describing SST bimodality and spell peaks (McClanahan et al., 2019, 2020a). The same reasoning applies to the modulation of heat stress by non-heat related environmental conditions (such as tidal range, cyclone frequency, human population density), which were found in previous works (Maina et al., 2011; Safaie et al., 2018; McClanahan et al., 2019, 2020a, 2020b; Sully et al., 2022) but that wasn’t the focus of the current study.

Finally, we used taxonomic groups primarily defined by taxonomic family and growth form and assumed that species belonging to each of these taxa had similar associations with environmental trends. In reality, this might not necessarily be the case, as it is possible that within these taxa there are species with divergent heat associations. Furthermore, it is important to underline that each heat association observed is an average for a given taxon, and that for this average we are not able to evaluate the specific weight of single species. Future work could focus on re-annotating the CSS surveys with more specific taxonomic labels (for instance, at the genus level), so that the analyses can be refined with a higher taxonomic resolution.

### 4.3 Perspectives

We observed that local differences in coral cover matched distinct patterns of interaction between recent and past heat stress depending on taxa. These distinct patterns could highlight differences in how taxa respond to past exposure to heat stress, and future work should investigate the possible causes.

As discussed above, one possible cause is a community shift from heat sensitive to heat tolerant species. To evaluate this hypothesis, future studies could focus on field survey data at higher taxonomic resolution than in our analysis, to allow heat associations to be assessed at a species level. Such studies are likely to be hampered by difficulties in defining coral species based on visual surveys (Postaire et al., 2016; Forsman et al., 2020; Oury et al., 2020; Prada and Hellberg, 2021; Rippe et al., 2021). This could be overcome by using molecular surveying techniques such as environmental DNA, which could make it possible to establish objective molecular boundaries to distinguish between surveyed species (Shinzato et al., 2018, 2021; Dugal et al., 2022).

These presumptive responses to past heat exposure could also be mirroring micro-evolutionary processes, where individuals with genetic traits conferring thermal tolerance are persisting after recurrent heat exposure. To investigate this, population genomic studies featuring genotype-environment association analyses could be performed to evaluate the emergence of adaptive genetic traits at reefs exposed to heat stress over the past decades (Riginos et al., 2016; Selmoni et al., 2020b, 2021).

Another possible mechanism underpinning these association with past heat is the physiological acclimation of corals exposed to recurrent thermal stress. This phenomenon could be verified by performing standardized and systematic eco-physiological analyses, which can be done using portable experimental systems to test heat tolerance (e.g. the Coral Bleaching Automated Stress System, CBASS; Voolstra et al., 2020).

In reality, it is likely that each of the three phenomena discussed here act together in concert to shape the heat response of a reef exposed to recurrent thermal stress. Understanding and disentangling the individual contribution of each phenomenon will be aided by running trans-disciplinary studies (i.e., performing ecological surveys, genomic and eco-physiological analyses in parallel) across the same reef system.

### 4.4 Conclusions

We performed an association study across reefs worldwide that revealed differences in local coral cover associated with heat stress trends. These associations differed between coral taxa, which we sorted into four groups accordingly:

**Group 1: Branching, corymbose, tabular and digitate Acroporidae.** Reefs exposed to recent heat stress displayed lower coral cover than locally expected; among these reefs, those previously exposed to frequent past heat stress showed relatively higher coral cover, when compared to those less frequently exposed.
**Group 2: Massive and encrusting Poritidae and meandroid Favidae and Mussidae.** Reefs exposed to recent heat stress displayed lower coral cover than locally expected, yet this association was weaker than for Group 1. No mitigating interaction of recent heat stress with past heat stress was observed.
**Group 3: Branching Pocilloporidae.** The spatial overlap between differences in local coral cover and recent heat stress was only observed when considering the interacting role of past heat stress. Indeed, reefs exposed to frequent past heat stress showed a positive association between recent heat stress and coral cover, while a negative association was observed when past heat stress was less frequent.
**Group 4: Branching Poritidae.** The associations between coral cover and heat stress trends appeared to follow a similar trend to Group 3, although this was weaker, noisier and not statistically significant.

The groupings presented here could represent coral taxa-specific response to heat stress exposure across reefs worldwide over the past two to three decades. Future studies will need to validate the existence of such processes and investigate their nature (community shifts, evolution or acclimation) in further depth and at higher resolutions.

## 5 Conflict of Interest

The authors declare that the research was conducted in the absence of any commercial or financial relationships that could be construed as a potential conflict of interest.

## 6 Acknowledgments

We thank the Catlin Seaview Survey project for collecting and giving access to the field survey data, the Coral Reef Watch for granting access to the degree heating week data, the United Nations Environment Programme World Conservation Monitoring Centre for granting access to the worldwide distribution of coral reefs data and the Coral Trait Database for granting access to the data on coral physiological, ecological and morphological traits. We thank Annie Guillaume and the anonymous reviewers for the comments and suggestions provided during the redaction of this paper.

## Notes

### Competing Interest Statement

The authors have declared no competing interest.

### Summary of Updates

Update after revisions

