## Supplementary Figure 1 for "Worldwide analysis of reef surveys sorts coral taxa by associations with recent and past heat stress"

### *Supplementary Material*

**Supplementary Figure 1. Correlation between environmental variables.** (A) Variables used to describe long-lasting environmental trends; (B) variables used to describe period-specific heat stress trends; (C) environmental variables describing recent heat stress ($DHW_{MAX\_1yr}$) and past heat stress over the long-term ($DHW_{...\_>1yr}$).

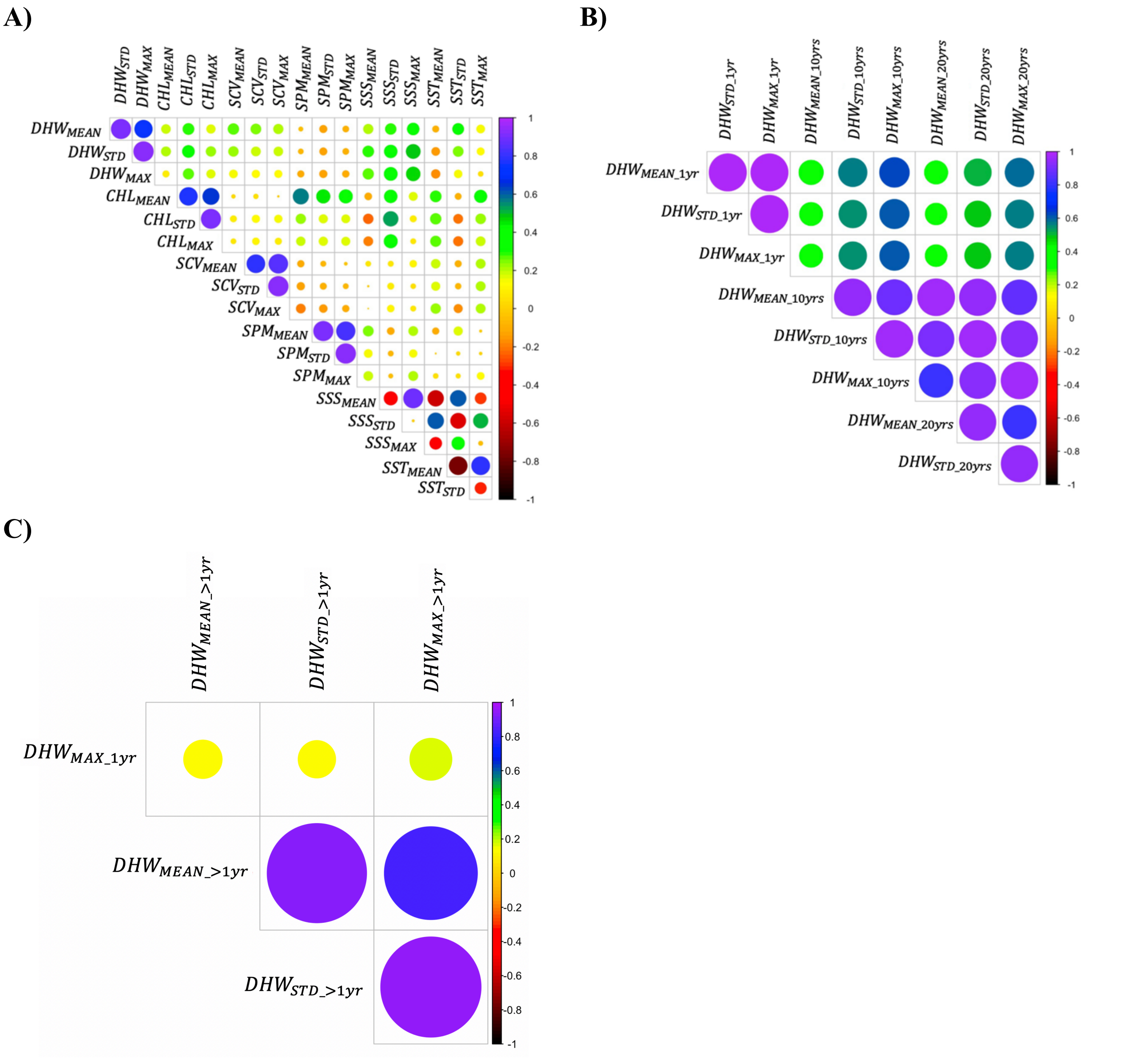
